# The abnormal, mis-localizated HR bmh protein associates with members of the protein processing machinery in the cytoplasm

**DOI:** 10.1101/245423

**Authors:** Eric G. Folco, Maud-Virginie Brancaz-Bouvier, Agnès Belly, Stefan Nonchev

**Author notes:** correspondence to: Stefan Nonchev Institut for Advanced Biosciences, INSERM 1209, UMR CNRS 5309, Université Grenoble-Alpes, Grenoble, France. Current Address: Université de Lyon, Institut de Génomique Fonctionnelle de Lyon, CNRS UMR 5242, Ecole normale supérieur de Lyon, Université Claude Bernard Lyon 1, 46 Allée d’Italie, F-69364 Lyon, France.

## Abstract

We have recently mapped a 296 bp deletion at the mouse hairless locus that causes the mutation hairless rhino bald Mill Hill, (*Hr^rhbmh^*), (ID:MGI:3039558; J:89321), removing the stop codon and generating a larger mutant protein HR bmh with an additional sequence of 117 amino acids. The mutant *hairless* gene mRNA is expressed during the embryonic and post-natal development of the hair follicle. A mutant HR protein was identified in bmh mouse skin at different stages of development by a specific antibody. We demonstrated that the HR bmh protein is able to interact with the vitamin D receptor, but is not able to repress VDR-mediated transactivation. Immunofluorescence analysis revealed that HR bmh protein displays an abnormal cellular localization in transfected cell lines, as well as in the epidermis and hair follicle of bmh mutant mice. Here we analyse the patterns of HR bmh extra-nuclear localization in cell transfection experiments. Using a candidate approach, double transfection experiments, immunofluorescent staining and IP protocols we established that HR bmh co-localizes specifically with the proteins HDAC6 in the cytoplasm is able to physically interact with it. Blast analysis allowed to show that HDAC6 and share high sequence homologies with specific motifs of HR bmh. We studied the association of these potential interactors with various cytoplasmic compartments. We discuss the relevance of the mutant hairless protein mis-localization in endosomal processing and in proteasome related pathways with respect to the specific skin phenotype of mouse hairless mutants.

## Introduction

In Mammals the gene *hairless* (*Hr*) encodes a nuclear factor strongly expressed in skin and crucial in controlling hair follicle integrity and cycling. In the absence of a normal and functional Hairless protein, the hair bulb undergoes premature apoptosis linked to loss of hair follicles and formation of epidermal utricles and dermal cysts (1), (2), (3).

The HR protein is localised in cell nuclei, tightly associated to nuclear matrix specific bodies and functions as a transcriptional regulator. Although its role has not been entirely resolved in molecular terms, it was demonstrated that HR is a corepressor for nuclear hormone receptors, HR is a component of large multiprotein complexes able to repress transcription by its association to chromatin remodelling factors such as histone deacetylases (4), (5), (6), (7). The *Hr* gene function is involved in cell adhesion modulation, *Hox* gene regulation, Wnt-signalling, and establishment of hair follicle progenitor cells identity (8), (9), (10). In the skin, as well as in other organs, the HR repressor seems to be a factor in the process of epithelial stem cells migration and differentiation. As the hair follicle undergoes repeated cycles of growth and degradation, these dynamic changes require stark and coordinated regulation of gene expression. The HR protein nuclear localisation is a key factor thought to control hair cycle dependant downstream gene transcription. A number of molecules involved in the control of hair cycling, not only change their activities and levels of their expression at the transition between anagen – catagen and catagen – telogen stages of hair cycles, but also modify their subcellular localisation according to the dynamics of growth and regression (11), (12). Shuttling between nucleus and cytoplasm has been reported for several proteins as a mechanism by which gene expression is regulated within different cell populations and components of the follicle itself. Mutated proteins, unable to fulfil their correct function can be directed to cytoplasmic compartments and processed for degradation. The class II histone deacetylase HDAC6 is a molecule that retains its capacity to acetylate histones and other proteins and in the same time it is predominantly localised in the cytoplasm (13). It has been shown that this enzyme has the intriguing property to be capable of nucleocytoplasmic shuttling (14). In some cases HDAC6 can translocate to the nucleus but its function in the cytosol remains elusive. This molecule has been shown to play a part in the ubiquitin signalling pathway and be involved in the proteasome dependant protein degradation (15). Unlike other nuclear histone deacetylases (HDAC1, HDAC3 and HDAC5), HDAC6 has not been found responsible for the HR protein corepresor function (6).

We have recently described a novel hairless mutation in the mouse – hairless rhino bald Mill Hill, symbol (*Hr^rhbmh^*) habouring a large 296 bp deletion at the 3’ part of exon 19 of the hairless gene (ID:MGI:3039558; J:89321), (9). We have shown that this deletion removes the stop codon, and creates a new reading frame of the hairless protein, generating a mutant product – HR bmh, encompassing an additional sequence of 117 amino acids at its C-terminal end (AS-117). This higher molecular weight protein is present in the embryo and at different stages of postnatal development. By contrast with its wild-type counterpart, HR bmh is mislocalised in the cytoplasm of transfected cell lines, epidermal and hair follicle cell populations (16).

Our present paper explores in more details the patterns of cytoplasmic localisation of HR bmh as well as its association with molecules involved in protein processing activities. We identify HDAC6 as a potential partner of the HR mutant protein.

## Results

### Nuclear versus cytoplasmic Hairless protein localization

To generate expression constructs and address the subcellular localization of the HR bmh protein we have previously cloned wild type and bmh mutant cDNAs in vectors tagged with the HA and the Flag epitopes. By single transfections in a variety of cell lines, we have shown that wild-type HR protein is nuclear, while the HR bmh mutant product is localized only in the cytoplasm and absent in the nuclei (16). As a first step to confirm and extend this analysis, we have now performed double transfections in Cos cells to coexpress mutant and wild type proteins in the same cell. In addition, in order to minimize artefacts of transfection, we swooped the tags HA and Flag of both proteins in the series of coexpression experiments. To detect the proteins we used Rhodamin coupled secondary anti-HA antibody and FITC-linked secondary anti-Flag antibody (see Material and Methods). Figure 1 A to F shows double transfection of Cos cells by HA tagged HR bmh and Flag tagged wild-type HR. In cells transfected with both plasmids (Figure 1, A, B, C and D) the wild type HR protein is nuclear (green), while the HR bmh (Red) is localised in the cytoplasm. In cells where only one plasmid was transfected, again as previously shown, the normal protein is nuclear and its mutant counterpart – cytoplasmic (Figure 1, E and F). Using the opposite labelling, i.e., when wild type HR is marked by HA and mutant HR by Flag, we note the same clear subcellular separation of the two proteins. (Figure 1, G to I). These data confirm that independently of the labels used, in cell culture experiments the HR bmh protein displays a steady and reproducible localisation in the cytoplasmic compartment. As the transfection efficiency vary from line to line, at least 20 to 30 fields were analysed for each double transfection experiment. A careful examination of double transfection and coexpression experiments confirmed our previous observations suggesting that in about 20% of the transfected cells; the immunoreactivity was not uniform, but rather concentrated in discrete cytoplasmic regions (16). We then set out to explore in more detail the way HR bmh is expressed in transfected cells.

**Figure 1.**
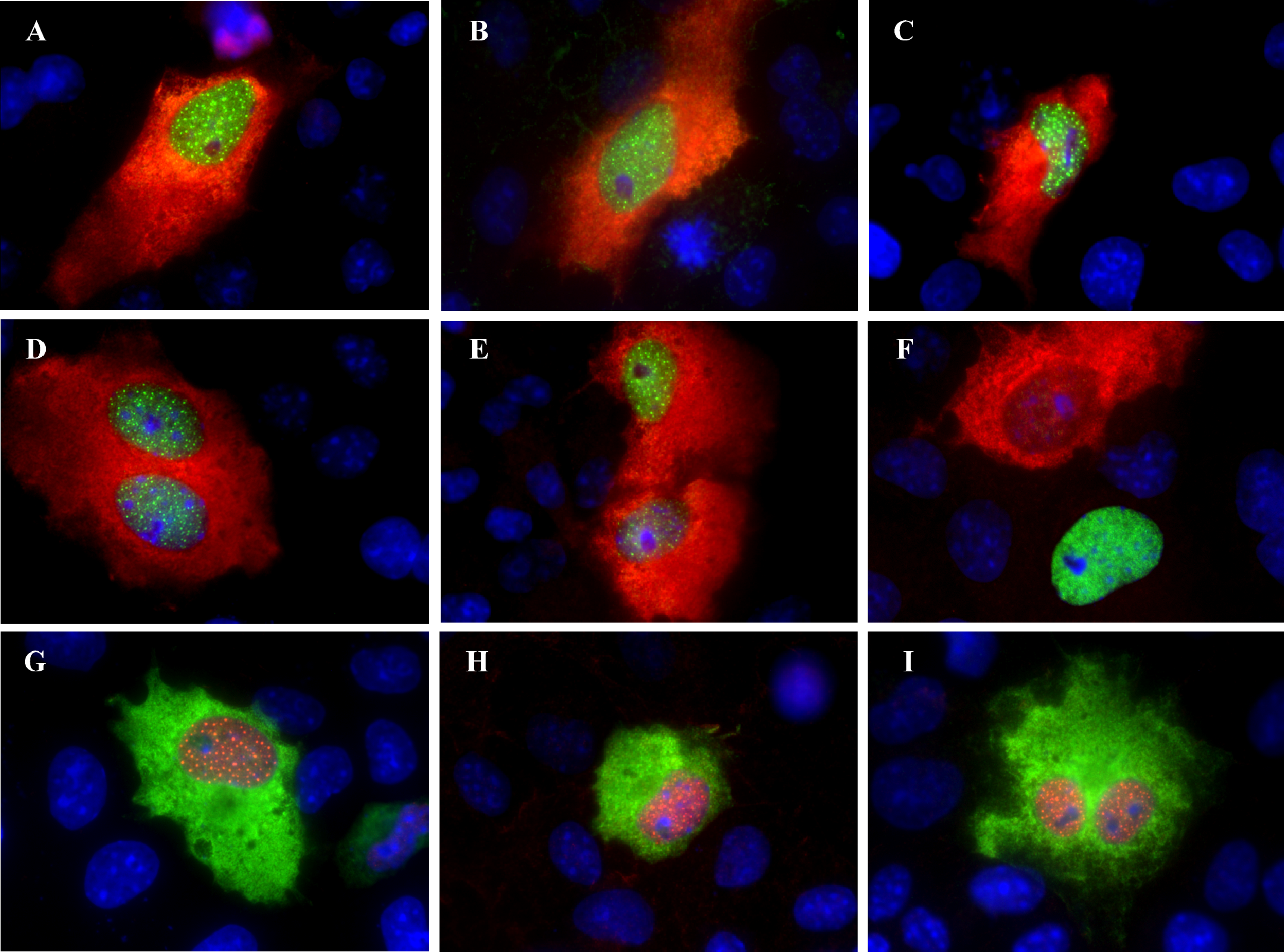
Coexpression of both wild-type and mutant HR in the same cell has no effect on their respective localisation. Cos cells were cotransfected with both Flag-Hr wt and HAHr bmh expression vectors. Subsequent localization of each protein was revealed by immunofluorescence using anti-Flag and anti-HA antibodies. As shown on panels A-F, the FITC-labelled (green) wild-type HR protein displays a nuclear localization, while rhodamine (red) labelled HR bmh remains cytoplasmic. On panels G to I, the wild type HR is marked by rhodamine in the nuclei, while HR bmh is FITC – stained in the cytoplasm.

### The HR bmh protein displays various patterns of cytoplasmic localization

The patterns of HR bmh irregular distribution are illustrated on Figure 2 A to I. The immunoreactivity is typically punctuated, localized in a number of compacts speckles with variable size either all over the cytoplasm, or confined in particular zones (Figure 2A, B and C). In these cells some areas are completely devoid of immunofluorescence (Figures 2A), while others display either filamentous or finely granulated staining (Figure 2B and C). The high variability in the repartition of the HR bmh product in the cytoplasmic compartment of the transfected cells is also characterised by juxtaposed zones of intensely stained cytoplasm surrounding areas where the protein is not expressed (Figures 2D, E and F) These “bmh-less” territories often display a marked perinuclear localisation. (Figure 2E, G and H). In striking contrast with this pattern, in many transfected cells the HR bmh protein is organised in dense cytoplasmic masses compactly apposed to the nuclear membrane with extremely highly levels of immunofluorescent staining (Figure 2, I). Although this odd distribution of the protein could be due to transfection and/or overexpression artefacts, it was to be taken cautiously into consideration while we were looking for cytoplasmic partners of the HR bmh protein.

**Figure 2.**
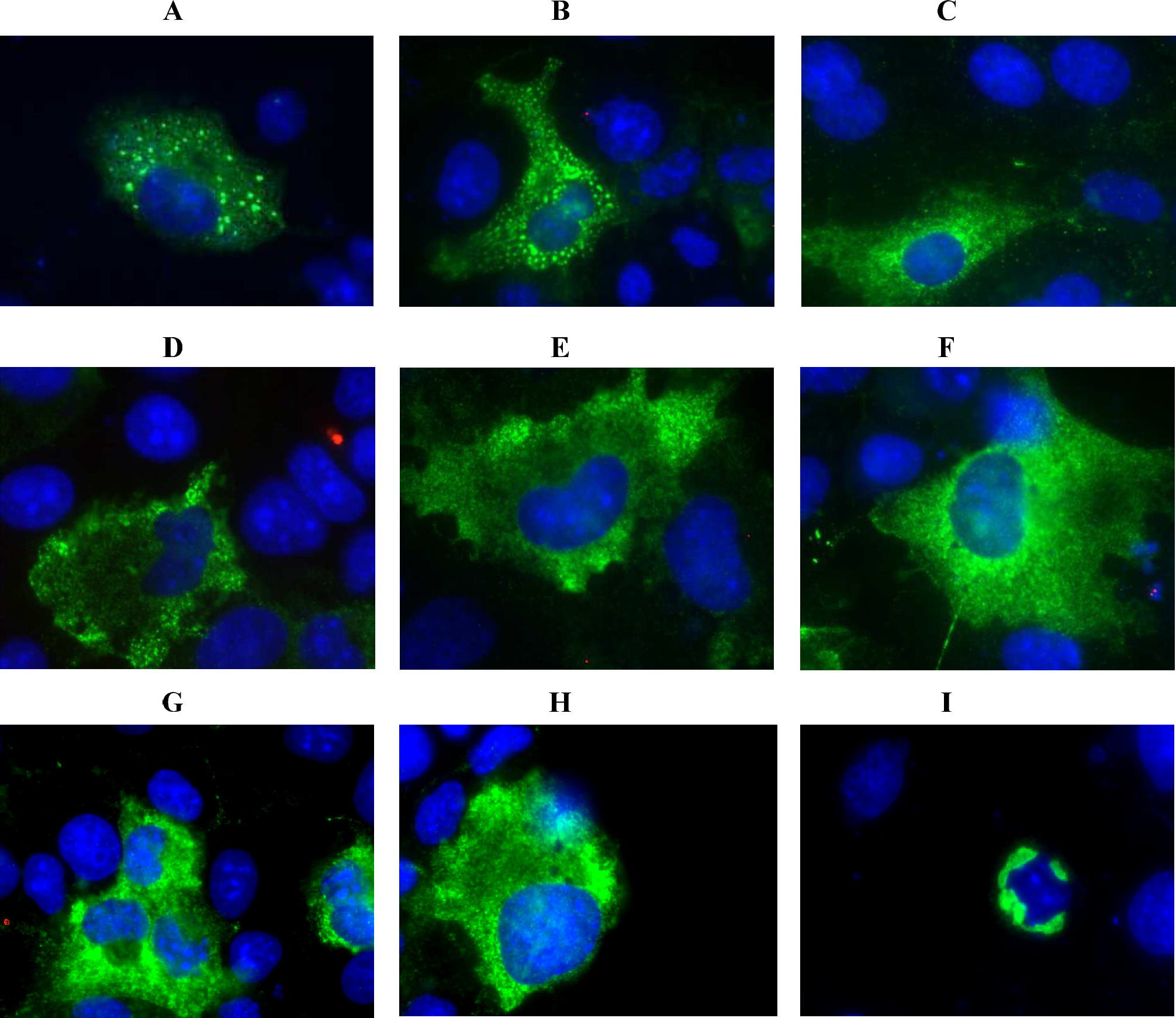
Patterns of cytoplasmic distribution of the HR mutant protein in transfected Cos cells. In addition to the homogeneous expression in the cytoplasm of transfected cells shown in Figure 1, the HR bmh protein is detected in perinuclear domains that can be highly condensed (A and I) or contain compact centres interspersed with specifically stained but decondensed cytoplasmic areas (B, C and F). The mutant protein can also be organised in small and compact speckles distributed all over the cytoplasmic area (F, G and H).

### The C’-AS117 putative NES is not functional in transfection experiments

As mentioned above and reported previously, the mutant HR bmh protein harbours an additional sequence of 117 amino acids at its C-terminal end (16). The rest of the protein is identical to the wild type one, in particular, the nuclear localisation signal (NLS) is intact, although its function is apparently perturbed (Figure 3A). To understand why this aberrant product is miss-localised in the cell, we set out to determine the precise role of AS117 in the behaviour of HR bmh. Our first approach was to verify if the addition of 117 amino acids could alter the correct tri dimensional folding of the wild type protein and hinder the functioning of the NLS. To tackle this possibility, we created a fusion protein with the GAL-4-DBD, known for its exclusive nuclear localisation. Transfections of the fusion protein GAL-4-DBD-HR bmh in a number of cell lines resulted again to clear cytoplasmic localisation for the fusion product (data not shown). These observations suggested that motifs responsible for cytoplasmic sequestration should be looked for in the 117 amino acid fragment. Careful examination of AS-117 allowed to spot the presence of a short sequence - LLLPLSSLSF- strongly resembling to a leucine rich consensus motif LxxxLxxLxL, called nuclear export signal (NES) and known for its capacity to regulate the nucleo-cytoplasmic transport of many proteins. To test the function, we targeted the C-terminal putative nuclear export signal motif of AS117 by specific deletions and mutations (Figure 3A). We created two variants of cDNAs coding for a HR bmh protein where the motif of interest is either mutated or deleted. By site directed mutagenesis we generated the construct HR bmh L83A/L87A, where the lysines 1 and 4 of the motif have been replaced by arginine. In the construct HR bmh CD35 the motif of interest as well as the last 25 C-terminal amino acids were deleted (Figure 3A). These two cDNAs were cloned in Flag-tagged expression vector and transfected in NIH 3T3 cells to analyse the effects of these mutations on the subcellular localisation of the HR bmh protein. As illustrated on Figure 3B, this experiment did not allow to define the precise element, responsible for the sequestration of the HR bmh protein. Neither the point mutation, nor the loss of the entire motif -LLLPLSSLSF- did modify the cytoplasmic localisation of the mutant Hairless protein. As the examination of the remaining N-terminal of the additional sequence does not contain putative sequences involved in a particular subcellular localisation, nor consensus sequences implicated in protein-protein interaction, we decided to perform a BLAST analysis in bank protein bank with C’-AS117 alone. Intriguingly, in the only hit of this BLAST, we found a strong homology between the central part of AS117 and a motif of the serum b2 microglobulin (Figure 3C), which is a non glycosylated single chain protein present in light chain of HLA-class I complexes, known to stabilise the tertiary structure of associated partners and play a role in the protein trafficking as well as a prognostic value marker in renal diseases. This finding incited us to explore a possible involvement of the mutant HR protein in various pathways of cytoplasmic of protein processing.

**Figure 3.**
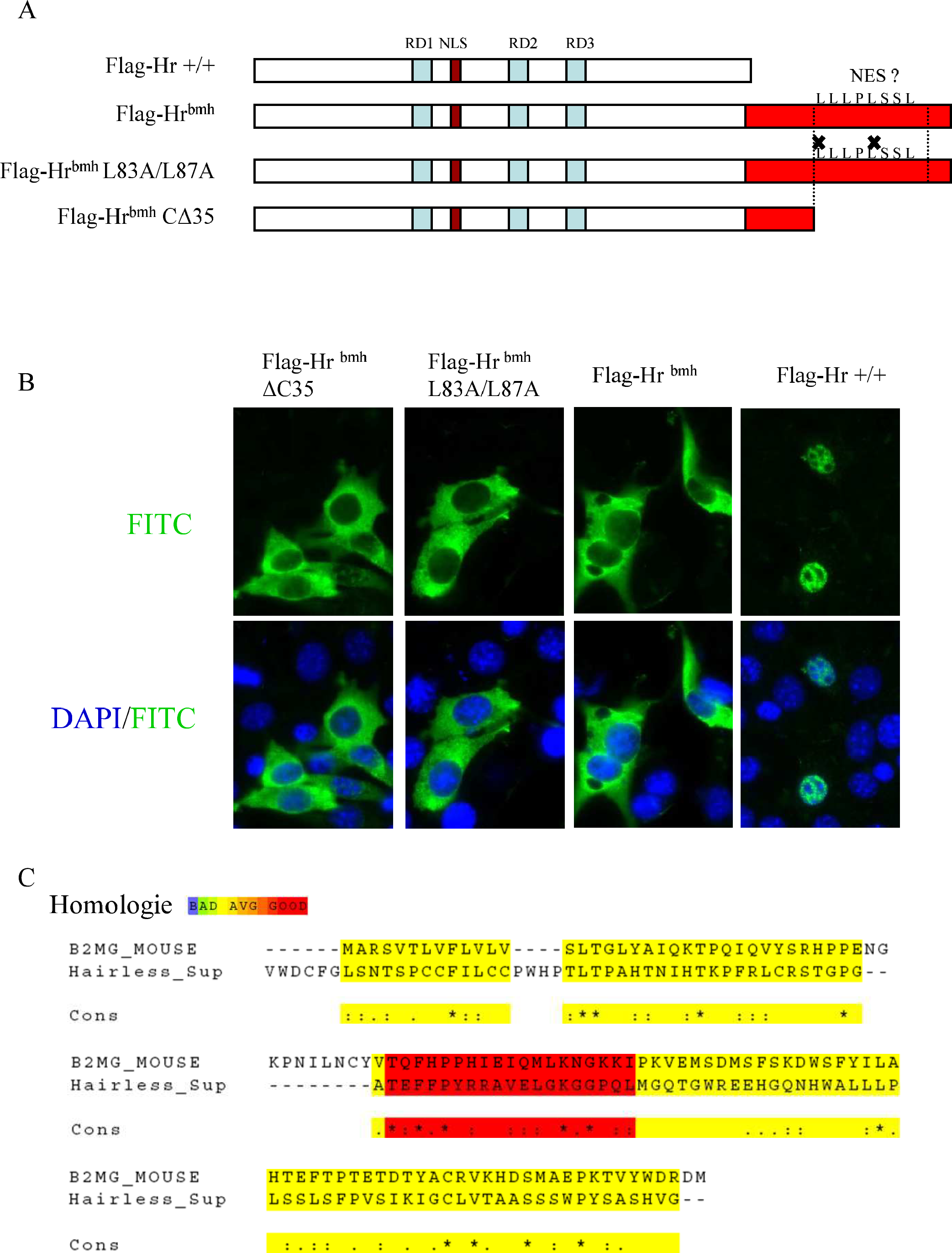
Mutation or deletion of the AS-117 of the HR bmh putative NES do not affect its cellular localisation. (A). A schematic representation of the fusion proteins Flag-HR wild type, Flag-HR bmh, Flag-HR bmh mutated at the putative NES motif (residus leucine 83 and Leucine 87) and Flag-HR bmh bearing a deletion of the putatif NES motif. (B). Indirect immunofluorescence using a anti-Flag antibody on NIH 3T3 cells transfected by constructs expressing the four proteins described in (A). Neither the point mutation (Flag-HR bmh L83A/L87A), nor the deletion (Flag-HR bmh CΔ35) of the putative NES motif affects the cytoplasmic localisation of these proteins. (C). The BLAST of the AS117 motif in protein banks reveals domains of high homology over about 30 amino acids between the central part of the AS117 and N-terminal and central domains of the small membrane antigen associated β2-microglobulin.

### The proteins HR bmh and HADC6 co-localise in the cytoplasm after transient transfection

The wild type hairless protein associates in vivo preferentially to histone deacetylases 1, 3, 5 and is thought to function as a corepressor through direct or indirect interactions with HDACs (4), (17). The HR bmh exerts its effects either by the loss of function as a nuclear corepressor, or by specific involvement in proteins trafficking and degradation in the cytoplasm. Looking for a cytosolic candidate able mediate protein degradation, we concentrated our analysis on the cytoplasmic class II histone HDAC6, which is implicated in acetylation of cytoplasmic proteins and proteasome processing (15). We co-transfected Cos cells with constructs expressing HR bmh and HDAC6 and analysed their patterns of expression after transient transfection. Intriguingly; the Cos cells expressing both proteins display highly reproducible patterns of immunoreactivity characterized by a number of specific features (Figure 4A to I). HR bmh and HDAC6 co-localise in large cytoplasmic areas, which are not uniformly stained, but present homogeneous zones interspersed with dots and speckles expressing high levels of both proteins (Figure 4A to I). Besides, HDAC6 specific fluorescence is observed in parts of the cytoplasm of variable size and location (Figure 4A to I). HR bmh also present zones where it is expressed alone, but these are of restricted size and most often apposed to the nuclear membrane (Figure 4A to I). The patchwork of overlapping and independently detected immunostaining raised the question of whether or not the HR bmh and HDAC6 can physically interact. To generate cytoplasmic extracts for immunoprecipitation experiments, we cotransfected NIH 3T3 cells with constructs expressing HA tagged Hairless wild type or mutant protein together with Flag epitope labelled HDAC6. The complexes were precipitated with anti-Flag antibody, the beads washed, and boiled before Western blot analysis with anti-HA antibody. As illustrated in Figure 4J, both wild type and bmh proteins immunoprecipitate with HDAC6 (Figure 4J, lanes 1 and 2). The specificity of this interaction was confirmed by the opposite combination of antibodies (data not shown) and a number of test transfections were used as controls (Figure 4J, lanes 3 to 7). These data were in favour of an involvement of HDAC6 in the processing of HR bmh and we wanted to know what part (s) of the HDAC6 molecule was implicated in the association and more importantly, which of its multiple functions was exposed to the interaction.

**Figure 4.**
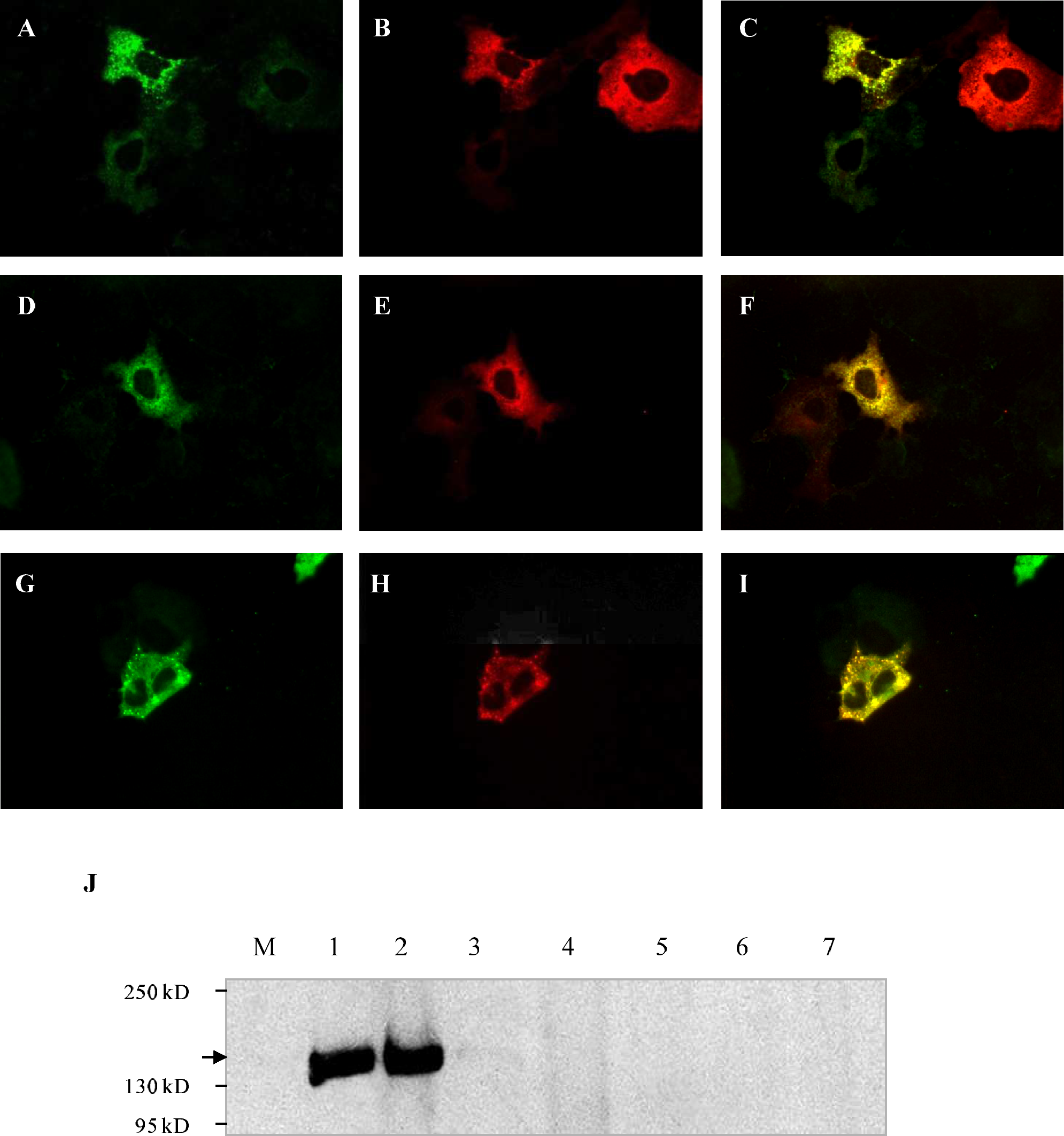
HR bmh and HDAC6 co-localize in the cytoplasm. A to I – Cells co-expressing Flag-tagged HR bmh (A, D and G, green) and HA-tagged HDAC6 (B, E and H, red). C, F and I show the merge of the two colours. The immunopecipitation assay (Materials and Methods) is shown on J: Lane 1 – HA-HDAC6/Flag-HR bmh; Lane 2 – HA-HDAC6/Flag-HR wild type; Lane 3 – HA-HDAC6/Flag-pSK alone; Lane 4 – Flag-HR bmh/HA-pSK; Lane 5 – Flag-pSK; Lane 6 – HA –pSK; Lane 7 – Flag HR wild type/pSK.

### The DD1 domain of HDAC6 is required for association with the HR bmh

As a candidate for interaction with HR bmh, the HDAC6 presents several functional motifs that might offer different options for the specific type of protein-protein interaction. A series of deletion derivatives has been previously created to map the functionalities and the cellular localisation of the HDAC6 enzyme (13). We decided to take advantage of these tools and co-express each one of them together with the whole of HR bmh protein in transfected Cos cells. Thus Flag-tagged HR bmh and HA-tagged HDAC6 deletion derivatives were used and the expression of transfected proteins was now monitored by confocal microscopy (Figure 5A to R). The patterns of full length protein colocalisation confirm the results obtained by wide field microscopy. Indeed, as previously observed, the dots and speckles of co-localised immunoreactivity are interspersed with areas and nodules where each of the partners is independently and strongly expressed. (Figures 5A to C). Constructs encompassing the N-terminal portion of the HDAC6 and the deacetylation domain one (DD1) also drove enzyme expression in the cytoplasm and created patterns of co-localisation (Figure 5D to F). When the DD1 was tested alone the profiles of immunoreactivity were again marked by high levels of yellow staining in the merge of the pictures generated by the two antibodies (Figure 5G to I). Interestingly, with this later construct the colocalisation of HR bmh and HDAC6 is even more widespread than when the whole HDAC6 was transfected (compare Figure 5I and F to Figure 4C, F and I). By contrast, in the absence of DD1 the two transfected proteins did not colocalise at all (Figure 5J to R). When the deacetylation domain 2 (DD2) is used, HDAC6 displays a restricted expression in the nucleus (Figure 5J to K). If the same DD2 is linked to the C-terminal domain the expression is very weak and diffuse all over the cell (Figure 5M to O). Similar patters of extinct HDAC6 expression and absence of any co-localisation was established if only the C-terminal portion is cotransfected with the HR bmh (Figure 5P to R). As the C-terminal end of HDAC6 harbours the site of homology and interaction with the ubiquitine binding complexes, this latest profile requires more caution interpreting the association of HR bmh and HDAC6 as a part of the proteasome dependant protein degradation pathway. Taken together the results of the deletion analysis suggested that the HDAC6 deacetylation domain one is able to interact independently with HR bmh and might be responsible for the observed co-localisation with HR bmh. Finally a BLAST analysis revealed that DD1 present striking homologies with a novel motif in the HR protein situated at the N-ter of the protein (Supplemental data). Our data with the cytosolic component HDAC6 left open the question if HR bmh could associate to membrane structures of the cytoplasm. The last aim of this analysis was to focus on its possible partnerships with specific ultrastructures.

**Figure 5.**
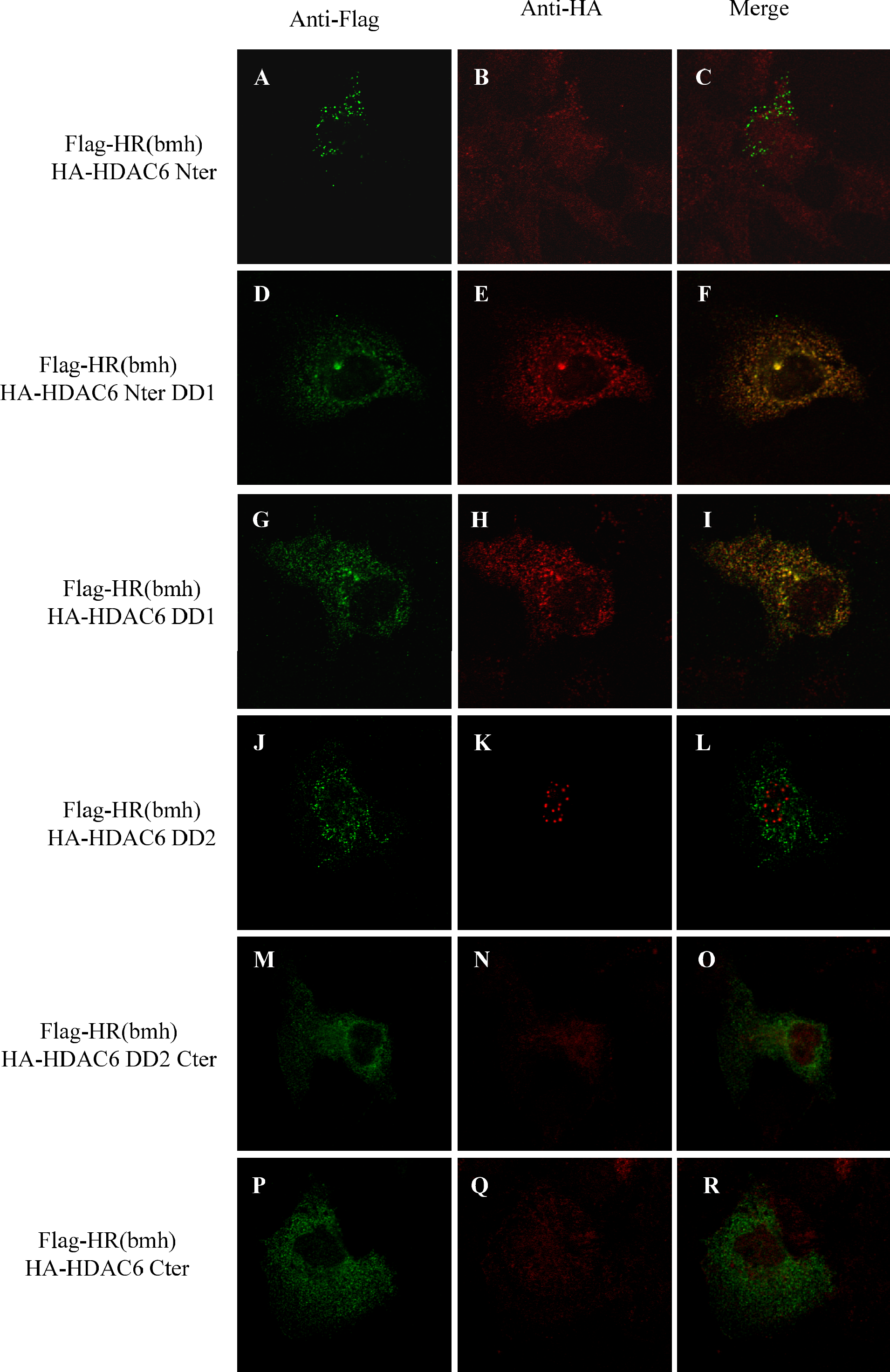
HDAC6 deacetylation domain 1 (DD1) is responsible for co-localisation with HR bmh. A to R illustrate co-transfections of Cos cells with Flag-tagged HR bmh and HA-tagged HDAC6 deletion derivatives. The pictures were generated by confocal microscopy. Note that the presence of the DD1 is associated with co-localisation of these two proteins (A to I). In constructs encompassing the DD1 domain alone the colocalisation of HR bmh and HDAC6 is even more widespread than when the whole HDAC6 was transfected (compare Figure 5I and F to Figure 4C, F and I). In the absence of the DD1 the proteins do not colocalise (J to R) and HA-HDAC6 displays either nuclear (K), or weak and diffuse distribution in all of cellular compartments.

### The HR bmh protein is detected in the lysosomal compartment

Bearing in mind the high homology of AS117 with the cell surface antigen b2 microglobulin, we wanted to establish if HR bmh could be found sequestrated in a specific cytoplasmic organelles. We took advantage of antibodies raised against Golgi apparatus (GM130), the endoplasmic reticulum (KDEL), the early endosomes (EEA1) and late endosomes and lysosomes (Lamp1). Cos cells were then transfected with Flag-tagged HR bmh protein and we analysed by immunofluorescence and confocal microscopy the mutant protein localisation with respect to the main subcellular compartments. Figure 6 A to F shows that HR bmh is not detected neither in the Golgi apparatus nor in the endoplasmic reticulum. Similarly Figure 6 G to L illustrates the absence of HR bmh in the early endosomes. By contrast, the antibody Lamp1, detecting late endosomes and lysosomes displays areas of co-localisation with the anti-HA tagged HR bmh (Figure 6M, N and O). These data indicate that HR bmh could be associated with the late endosomes/lysosome compartment. As a whole our analysis is in favour of the idea that HR bmh could be either taken by HDAC6 for cytosol associated processing, or follow a pathway of lysosomal degradation.

**Figure 6.**
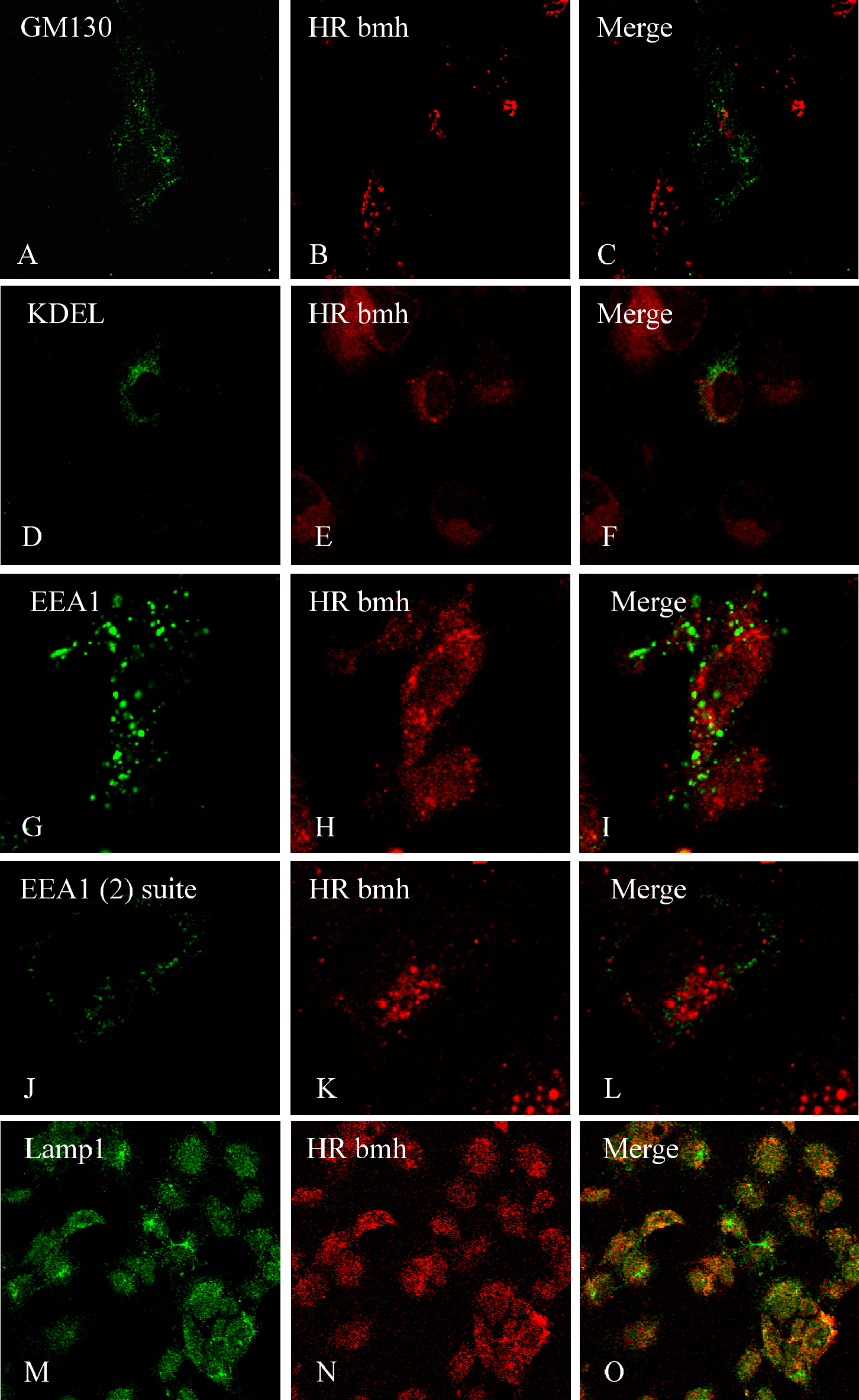
HR bmh is associated with the late endosomes/lysosome compartment. Cos cells were transfected with HA-tagged HR bmh and antibodies raised against the principal cytoplasmic ultrastructures were used to detect organelles where HR bmh is located. The pictures were taken by confocal microscopy. A, B and C show that HR bmh is not found in the Golgi apparatus. D, E and F suggest that the endoplasmic reticulum does not associate with the Hairless mutant protein, while G to L illustrates its absence in the early endosomes. Note that the antibody Lamp1, detecting late endosomes and lysosomes displays areas of co-localisation with the anti-HA tagged HR bmh (M, N and O).

These options are to be further explored with appropriate reagents and experimental approaches.

## Discussion

In this work we have investigated in cell culture experiments several aspects of the cytoplasmic sequestration of a mutant protein, thought to underlie the mouse bald Mill Hill hairless phenotype (9), (16). We show that HR bmh displays a heterogeneous distribution in the cytoplasm of transfected cells, either in aggregates of nodules, speckles and fibres, interspersed with zones devoid of the protein, or in highly compact areas tightly apposed to the nuclear envelope. We tested the role of the additional sequence AS117 that characterizes the Hairless bmh protein by mutating and deleting the putative LXXLL nuclear export sequence to find out that this part of the protein was not involved in the cytoplasmic localisation. The cytosolic enzyme HDAC6 was identified as a potential partner of HR bmh. Indeed, the IP assays demonstrate that these proteins are able to bind to each other and they co-localise in distinct cytoplasmic zones. Finally, based on homologies between the AS117 fragments and the small cell-surface antigen β2 microglobulin, we explored the possible link between HR bmh and the membrane ultrastructures and established that the mutant protein is likely to associate with the late endosome/lysosome compartment.

The heterogeneous distribution of HR bmh is something we treat with extreme caution. Indeed, overexpression of a nuclear protein by transient transfection, especially in Cos cells may be also a transfection artefact. Without making a major point about that, we still consider it as a sign to look for compartments, molecules and functions that might implicate more specifically HR bmh. As the appropriate subcellular localisation is crucial for correct function, we wanted to take into account every possible cue that might point to a specific regulatory pathway. Signal induced activation of transcription factors that change their compartments of residence is well described in immune and inflammatory responses (18)(18), (19). Shuttling between nucleus and cytoplasm was reported for a number of transcription factors involved in the control of hair cycling. The vitamin D receptor, known to regulate the course of hair growth in anagen must be localised in the nucleus and then bind to the enhancers of hair cycle involved downstream genes (20), (16). At other stages of the hair cycle or in the absence of its ligand 1,25(OH)2D3 in cell culture experiments, the VDR is detected in the cytoplasm (21). Immunohistochemistry data have demonstrated that a number of other factors that display dynamic changes in their expression patterns in epidermis and hair follicle, might also present distinct differences in the subcellular localisation between hair follicle compartments. Thus, in the case of the C/EBP family of transcription factors shuttling between nucleus and cytoplasm is a mechanism to regulate activities in distinct cells populations within the hair follicle (12). Interestingly, the HR wild type protein, which has a dynamic pattern of expression in the progression of the hair cycle with highest levels during catagen and complete absence in anagen, has never been detected in the cytoplasm (8). Modulations in the localisation of transcriptional co-regulators relay on the presence of signature motifs like LXXLL, that might not be involved in direct interaction, bur are absolutely required for repressive or activating function (22). The LXXLL, originally identified in several nuclear receptor co-activators, is a protein interaction module that has been shown to mediate a variety of ligand induced responses. As the main function of this motif is thought to be linked with assembly of correpressor complexes in the cytoplasm, it is considered to be a part of an effective NES, although nuclear proteins might also harbour LXXLL-like motifs (23). In our hands neither the mutation, nor the deletion of the putative AS117 NES motif modified HR bmh cytoplasmic sequestration. Nevertheless, the AS117 sequence was shown to present homology with the small molecule b2 microglobulin, a common subunit of cell-surface antigens and involved in protein trafficking (24). This finding urged us to explore a cytoplasmic pathway for HR bmh, where the mutant protein could be a part of cell membrane compartments. Using specific antibodies we have shown that the late endosomes and the lysosomes could shelter HR bmh, while the endoplasmic reticulum, the Golgi apparatus and the early endosomes are not likely to be preferred sites for its sequestration. One way to explain this result is to consider that HR bmh would be recognized as an aberrant product by effectors of the lysosome pathway and tethered to the structures responsible for redirection for lysosome processing and degradation. A more tempting scenario is offered by the interaction of HR bmh with the cytosolic protein HDAC6. This enzyme is a regulatory agent involved in protein ubiquitination and proteasome dependant protein degradation (14,15,25). If such a partnership is real, the HR bmh could be a part of a Ub-associated signalling pathway, where its functions are not clearly predictable. The role of HDAC6 in this interaction is not likely to be just the one of a simple carrier of HR bmh to the proteasome; In fact, in our hands, the C-terminal, known to mediate binding to Ub is not involved in HR bmh – HDAC6 co-localisation. On the other hand, the C-terminal HR JmJ motif enzymatic functions have never been tested in the contexte of the additional 117 amino acid sequences lying immediately to the JmJ domain. In an alternative scenario, the important role of the DD1 domain of HDAC6 and its high homology with a novel functional motif at the N-terminal end of the HR protein is in favour of another, and yet unidentified type of partnership between the two molecules that can be dependant on their conformation, and is likely to involve their enzymatic activities and possible common substrates. Further work is required in order to establish an authentic link between the subcellular localisation of the HR bmh protein and the skin phenotype of the mutant hairless bald Mill Hill mice.

## Material and Methods

### Cell Culture and Transient transfection assay

Cell lines were purchased from the American Type Culture Collection (ATCC) (Rockville, MD). Media and reagents for cell culture were purchased from Gibco (Invitrogen) unless otherwise indicated. COS-7 cell lines was maintained in DMEM high glucose (4,5g/L) medium containing L-glutamine, 10% fetal bovine serum, 100 units/ml penicillin, and 100 μg/ml streptomycin at 37 °C under 5% CO2. Cells were plated on 60 mm dishes, each dish containing 1 × 10^6^ cells. Following a 24 h incubation, cells in each well were transfected with 5 μg of total DNA contruct using the SuperFect Transfection Reagent according to the manufacturer’s instructions (QiaGen). After 12 h of transfection, the cells were washed with phosphate-buffered saline and then used for Western blot or immunofluorescence experiments.

### DNA Constructs

Full length wild-type *Hr* and bmh *Hr* cDNA were subcloned into pcDNA3-Flag expression vectors as described previously (16). HA-HDAC6 and HDAC6 expression vectors were previously described (13). The sequences of all clones were confirmed by sequencing.

### Blast analysis

Hairless and HDAC6 proteins sequences were taken from Swiss-Prot database (accession number: Q61645 and Q9Z2V5 respectively). We added to the sequence the 117 amino acids of bmh mutant form as previously described (16). The blast analysis was processed on T-Coffee (Version 5.13 from Swiss Institute of Bioinformatic) (26).

### Immunofluorescence procedures

Cells were plated onto a 2 wells (2×10^5^ cells/well) labtek chamber slide (Nalgen Nunc International). Cells were transiently transfected with plasmid constructs as described above. Two days after transfection, cells were fixed in 4% paraformaldehyde (PFA) in phosphatebuffered saline (PBS) for 5 min at room temperature and permeabilised by the addition of 0.1% Triton X-100. Incubation with primary antibodies (refer to antibodies section) was carried out overnight at 4°C in the PBS-milk solution. After the addition of the secondary antibodies, cells were washed and counterstained with Hoechst 33258. The slices were then observed under a wild field microscope (Zeiss Axiophot) and confocal microscope.

### Immunoprecipitation and Western blotting procedures

200 μg of a cytosoluble cell extract were incubated with Anti-Flag M2 coupled beads (Sigma) for 2 h. After washing, beads were boiled in loading buffer and analyzed by Western blot: The beads were separated by SDS-PAGE followed by transfer onto a nitrocellulose membrane. The membranes were blocked with 5% non fat dried milk in TBST (20 mM Tris-HCl (pH 7.5), 500 mM NaCl and 0.05% Tween 20) and incubated for 2 hours with primary antibody (mouse monoclonal anti-HA(11)). The membranes were washed three times for 10 min each with TBST and then probed with secondary antibody (anti-mouse coupled to HRP), washed again with TBST, and developed with ECL Western blotting detection system (Pharmacia Biotech Inc.). Anti-phosphatase cocktail 1, B-Glycerophosphate (Sigma G6251; 25 mM), NaF (Sigma S7920; 5 mM), Complete Tabs (Roche), 50mM MgCl2, and 5mM CaCl2 were used during each step of immunoprecipitation.

### Antibodies

For immunoprecipitation we used as primary antibody:

– The mouse monoclonal anti-HA(11) (Reference: F3165 from Covance)

As secondary antibodies:
– The polyclonal goat anti-mouse coupled to HRP (Reference: P0447 from DAKO)

For Immunofluorescence we used as primary antibodies:
– The rabbit polyclonal anti-HA(Y11) (Reference: SC 805 from Tebu)
– The mouse monoclonal M2 anti-Flag (Reference: F3165 from Sigma)
– The mouse monoclonal anti-KDEL (Reference: SPA-827 from Stressgen)
– The rabbit polyclonal anti-EEA1 (Reference: ab29000 from Abcam)
– The rabbit polyclonal anti-LAMP1 (Reference: ab24170 from Abcam)
– The mouse monoclonal anti-GM130 (Reference 610822 from BD Transduction Laboratories). The KDEL, EEA1, GM130 and LAMP1 antibodies were a kind gift from Drs Agnès Belly and Yves Goldberg.

As secondary antibodies:
– The Alexa Fluor 488 goat anti-mouse (Reference: A 11029 from Invitrogen)
– The Alexa Fluor 546 goat anti-mouse (Reference: A 11030 from Invitrogen)
– The Alexa Fluor 488 goat anti-rabbit (Reference: A 11034 from Invitrogen)
– The Alexa Fluor 546 goat anti-rabbit (Reference: A 11035 from Invitrogen)

## Acknowledgements

We thank Drs Yves Goldberg, Cyril Boyaut and Saadi Khochbin for the generous gifts of reagents and helpful discussions. This work was supported by the “Emergence” grant of the Region Rhône – Alpes and by the French Fondation de la Recherche Médicale (SN). M-VB and EF had a PhD fellowship of the French Ministry of National Education. We are grateful to Brigitte Peyrusse for skilful artwork.

